# Abundance, density, and social structure of African forest elephants (*Loxodonta cyclotis*) in a human-modified landscape in southwestern Gabon

**DOI:** 10.1101/827188

**Authors:** Colin M. Brand, Mireille B. Johnson, Lillian D. Parker, Jesús E. Maldonado, Lisa Korte, Hadrien Vanthomme, Alfonso Alonso, Maria Jose Ruiz-Lopez, Caitlin P. Wells, Nelson Ting

**Author notes:** Corresponding Author: Nelson Ting, 308 Condon Hall, 1218 University of Oregon, Eugene, OR 97403. These authors contributed equally to this work (Colin M. Brand, Mireille B. Johnson).

## Abstract

The noninvasive monitoring of population size and demography is critical to effective conservation, but forest living taxa can be difficult to directly observe due to elusiveness and/or inaccessible habitat. This has been true of African forest elephants (*Loxodonta cyclotis*), for which we have limited information regarding population size and social behavior despite their threatened conservation status. In this study, we estimated demographic parameters focusing specifically on population size and density using genetic capture-recapture of forest elephants in the southern Industrial Corridor of the Gamba Complex of Protected Areas, which is considered a global stronghold for forest elephants in southwestern Gabon. Additionally, we examined forest elephant sociality through analysis of social networks, predicting that we would find matrilineal structure as exhibited by savanna elephants and other forest elephants. Given 95% confidence intervals, we estimate the size of the population in the sampled area to be between 754 and 1,502 individuals and our best density estimate ranges from 0.47 to 0.80 elephants per km^2^. When extrapolated across the entire Industrial Corridor, this estimate suggests an elephant population size of 3,033 to 6,043 in this area based on abundance or 1,684 to 2,832 based on density, which is 40 – 83% smaller than previously suggested. Furthermore, our social network analysis revealed approximately half of network components included females with different mitochondrial haplotypes; this suggests a wider range of variation in forest elephant sociality than has previously been reported. This study emphasizes the threatened status of forest elephants and demonstrates the need to further refine baseline estimates of population size and knowledge on social behavior in this taxon, both of which will aid in determining how population dynamics in this keystone species may be changing through time in relation to increasing conservation threats.

## Introduction

Compared to African savanna elephants, the biology and behavior of African forest elephants (*Loxodonta cyclotis*) remain poorly understood. This knowledge gap may be attributed to their disputed status as a distinct species from *L. africana* (Roca et al. 2001), their cryptic nature, and occupation of difficult to access habitat. Indeed, this last factor limits most behavioral observations of this species to time spent in bais (i.e. forest clearings around a water resource; Fishlock and Lee 2013, Querouil et al. 1999, Turkalo and Fay 1995), although observations have been possible when the elephants occupy coastal or savanna habitats (Morgan and Lee 2007, Mormont 2007, Schuttler et al. 2014b, White et al. 1993). Studies of African forest elephants are critical given their conservation status and recently documented population declines (Maisels et al. 2013, Poulsen et al. 2017), as well as their relevance for understanding the evolution of proboscideans (Meyer et al. 2017, Palkopoulou et al. 2018).

Whereas many studies have focused on the feeding ecology of this species (Short 1981, Tchamba and Seme 1993, White et al. 1993), fewer have examined their social behavior. Early models of forest elephant sociality assumed similarities with African savanna elephants, which exhibit a fission-fusion social structure with subunits consisting largely of multiple related females while males are primarily independent and do not consistently associate with a particular subunit (Archie et al. 2006, Douglas-Hamilton 1972, Moss and Poole 1983). Among savanna elephants, Wittemyer et al. (2005) describe a multi-tiered social structure with mother-calf pairs forming the simplest unit, families consisting of multiple mother-calf pairs, kinship groups being composed of multiple families, and clans consisting of multiple kinship groups. Direct observation of forest elephants has revealed that social groups are considerably smaller than those of African savanna elephants with groups typically composed of two to three individuals, often mothers and their dependent offspring (Morgan and Lee 2007, Turkalo et al. 2013). Genetic studies have confirmed that individuals in these small groups are usually highly related (Munshi-South 2011, Schuttler et al. 2014a, Schuttler et al. 2014b). Studies of forest elephants in bais have also revealed that while forest elephant groups differ in size compared to savanna elephant groups, both species exhibit fission-fusion social structure (Fishlock and Lee 2013, Schuttler et al. 2014b). Collectively, these data have furthered our understanding of forest elephant sociality; however, many of these studies have relied on observations of forest elephants in bais. Further study of forest elephants outside of bais are of great importance, especially considering that behavior may differ between bais and other habitats.

Many studies of forest elephants have been conducted in national parks or other protected areas (e.g. Fishlock et al. 2008, Munshi-South 2011, Schuttler et al. 2014a,b). However, forest elephants, like many other species, are experiencing increasing contact with humans outside of protected areas as a result of agricultural expansion, resource extraction, and increased urbanization occurring across sub-Saharan Africa and elsewhere. Assessing the potential effects of anthropogenic activity on wildlife biology and behavior by studying forest elephant populations in human-dominated landscapes is now a key research priority (Aherling et al. 2012, Ahlering et al. 2013, Johnson et al. 2019).

Forest elephant populations have been dramatically decreasing over the past few decades for a number of reasons including habitat loss and poaching (Blake et al. 2007, Maisels et al. 2013). One recent study highlighted major population loss (78-81%) over a decade within a protected area in northeastern Gabon (Poulsen et al. 2017). The authors argued that the primary reason for this decline is attributed to poaching. Collectively, these studies highlight the immediate need not only for measures to protect forest elephants but also for assessing baseline levels and changes in population size, density, and social structure. While genetic studies can be less cost effective than other methods, noninvasive sampling provides more accurate and precise estimates than other traditional census methods (e.g. Arrendal et al. 2007, Guschanski et al. 2009). Such census methods (e.g., counting dung piles) have provided initial global estimates of ∼100,000 remaining forest elephants (Blanc et al. 2007), with Gabon identified as containing a considerable proportion of individuals. In particular, the Gamba Complex of Protected Areas in southwestern Gabon, which consists of two national parks (Loango and Moukalaba-Doudou) and an intervening Industrial Corridor, has been described as a bastion for forest elephants. Thibault et al. (2001) estimated that the forest elephant population in the entire complex ranged from 10,236 to 12,174 in 1999. A recent study by Eggert et al. (2014) speculated that the Industrial Corridor of the Gamba Complex contained approximately 10,000 forest elephants in 2004, which would have been approximately one-tenth of the world’s remaining forest elephant population at that time (Maisels et al. 2013). In addition to information about estimated population size, Eggert et al. (2014) found that this corridor has a resident forest elephant population and also serves as additional habitat to elephants whose primary ranges are in the neighboring parks. Elephants in the Industrial Corridor were also found to occur in small, matrilineal groups (Munshi-South 2011) and exhibit lower fecal glucocorticoid concentrations than elephants inhabiting neighboring Loango National Park (Munshi-South et al. 2008).

The present study has two research questions. 1) How many forest elephants occupy the 891 km^2^ area surrounding the town of Gamba in the Industrial Corridor? Based on the previous population size estimate for the entire Industrial Corridor, we optimistically predicted that our study area would contain approximately 3,330 forest elephants assuming relatively homogenous densities across the landscape and a stable population size since 2004 (Eggert et al. 2014). We used genetic capture-recapture to estimate the abundance and density of elephants within our study area, and we also extrapolated these estimates to infer the total elephant population size across the Gamba Complex Industrial Corridor. 2) What is the social structure of these forest elephants? The social structure of African savanna elephant clusters around matrilines with a multi-tiered system (Archie et al. 2006, Moss and Poole 1983) and the limited amount of data on forest elephants appears to suggest this is also true for this taxon (Fishlock and Lee 2013, Munshi-South 2011, Schuttler et al. 2014). We predicted that social network analysis and genetic data would yield networks that are nearly exclusively composed of females with the same mitochondrial haplotype.

## Methods

### Study Area

The Gamba Complex of Protected Areas in southwestern Gabon (Figure 1) covers approximately 9,600 km^2^ and includes two national parks (Loango National Park, 1,550 km^2^; Moukalaba-Doudou National Park, 4,500 km^2^) in addition to an intervening Industrial Corridor (3,585 km^2^), which holds oil and timber concessions (Alonso et al. 2014, Eggert et al. 2014, Munshi-South 2011, Munshi-South et al. 2008). Our study was conducted within the Industrial Corridor in an 891 km^2^ landscape between Sette Cama in the north and Mayonami in the south (Johnson et al. 2019). This landscape consists of a variety of habitats including grassland, primary forest, secondary forest, and wetlands. The study area also has the highest human population densities in the Gamba Complex, ranging from < 1 to > 60 people per km^2^ (Vanthomme et al. 2013), and features multiple plantations, settlements, and the town of Gamba with a population of > 9,000 people (Alonso et al. 2014, Johnson et al. 2019). The development of this town and the surrounding area is largely the result of oil extraction, which began in the 1960s (Alonso et al. 2014, Vanthomme et al. 2013). Further, this landscape includes the infrastructure necessary for oil extraction including buildings, wells, and roads (Johnson et al. 2019). Thus, much of the study area is dominated by human residence and/or activity.

**Figure 1.**
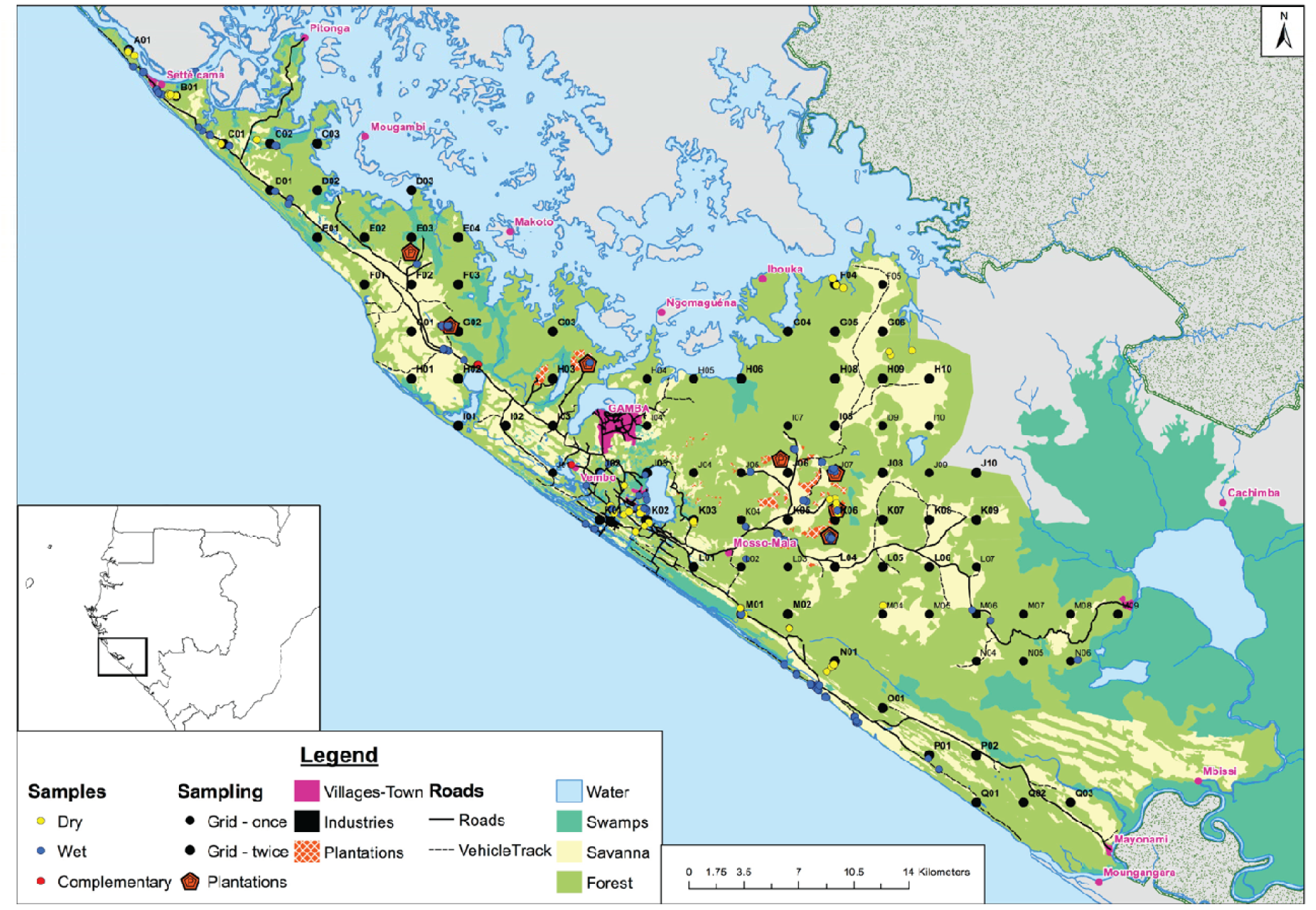
Location of study area, sampling grid, and sample localities.

### Sample Collection and Genotyping

Samples were collected non-invasively from fresh dung between June and August 2013 (during the dry season) and between October 2013 and March 2014 (the wet season). A sampling grid was constructed using ArcGIS 10.1 (ESRI, 2012) consisting of 100 points with 3 km intervals that covered the research area (Figure 1). On sampling days, three to four points were chosen at random. A team of three people searched for fresh dung 500 m before the point and 500 m after reaching the point in the direction of any elephant trails. Fresh trails from the point were also followed for ∼150 m.

Details on STR (short-tandem repeat) genotyping of dung samples are described in Johnson et al. (2019). Briefly, total genomic DNA was extracted from each sample for genetic analyses. Samples were genotyped at eight dinucleotide microsatellite (STR) loci (Supplement 1) and sexed using two Y-specific loci (*SRY* and *AMELY*) and one X-specific locus (*PLP1*) in a multiplex PCR reaction following Ahlering et al. (2011). We also sequenced a 600 bp segment of the mtDNA control region for all samples using previously established primers that amplify mtDNA in this taxon, MDL3 and MDL5 (Fernando et al. 2000).

GENALEX version 6.5 (Peakall and Smouse 2012) was used to determine P(ID), the probability that two individuals randomly drawn from a population have the same multi-locus genotype, and P(ID)_sib_, the probability that siblings have the same multi-locus genotype, which corrects for shared alleles among relatives and siblings. Thus, these measures test the power of a microsatellite panel to identify unique individuals (Waits et al. 2001). As elephants are highly social and may be associating with kin, we chose the more conservative approach: P(ID)_sib_, using a cutoff of 0.01. We used GENEPOP version 4.2 (Raymond and Rousset 1995) to test for deviations from Hardy-Weinberg equilibrium (HWE) and linkage disequilibrium (LD) across all markers. Fisher’s exact test was used to test for a deviation from HWE across all loci. We used the program’s Markov chain algorithm to determine departures from HWE and LD for each marker using 100 batches and 1000 iterations. Significance levels were adjusted using a Bonferroni correction. MICROCHECKER version 2.2.1 (Van Oosterhout et al. 2004) was used to detect the presence of null alleles. We used GIMLET version 1.3.3 (Valière 2002) to determine allelic dropout and false allele rates across all loci. GENALEX was used to identify unique individuals from the consensus genotypes, detect individual recapture, and estimate standard measures of genetic diversity. We considered samples with identical mtDNA haplotypes, sex, and STR genotypes to represent recapture of the same individual. We also allowed for samples to differ at one STR locus to represent recapture to account for genotyping error.

### Estimation of Abundance and Density

We used the package Capwire (Miller et al. 2005, Pennell et al. 2013) in R (R Core Team 2016) to estimate elephant abundance in our study area. Capwire generates abundance estimates from capture class frequency data. We classed individuals based on the number of times each unique individual was sampled. We considered samples to be recaptures if they were collected on different days. Capwire features three models to estimate population size: the equal capture model (ECM), the two-innate rates model (TIRM), and the partitioned two-innate rates model (TIRMpart). ECM assumes that all individuals in a given population have an equal probability of being sampled, whereas the TIRM categorizes each individual into one of two classes: individuals that are less likely to be resampled and those that more likely to be resampled (Miller et al. 2005). TIRMpart partitions the data into three different classes and calculates population size using TIRM, but excludes individuals from the third class that were detected a large number of times (Pennell et al. 2013). These individuals violate the assumption of two detection probabilities and may bias the population size estimate, and they thus are excluded. The number of individuals sampled in this third class are added to the point estimate calculated from the two classes for the final partitioned point estimate (Pennell et al. 2013). When generating point estimates we used a maximum population of 10,000 individuals. In addition to the point estimates from both models, we calculated 95% confidence intervals for each using parametric bootstrapping (N=100 bootstraps). We emphasize these confidence intervals, rather than the point estimates, as they are more informative for size estimation and monitoring trends over time (Arandjelovic and Vigilant 2018). All three models assume the target population is closed (e.g. no births, no deaths, no emigration/immigration). We also compared the ECM and TIRM as well as TIRM to TIRMpart using likelihood ratio tests with 100 bootstraps to determine which model best fit our data.

We also examined forest elephant density using the spatially explicit R package SECR (Borchers and Efford 2008, Efford 2011). This program estimates density by assuming each individual sampled has a circular home range and that capture probability decreases with increasing distance from the center of that individual’s home range. We used an exponential detection function, the default Poisson distribution of home range centers, and a buffer of 2000 m, which defines an area outside of the sampling area in which individuals may have a home range center. As some samples were collected near bodies of water (the Atlantic Ocean or the Ngodo Lagoon), we used a 2000 m buffer so as to not include home range centers in unsuitable habitat. This choice in buffer distance is further buttressed by known forest elephant home ranges from radio-collared individuals whose mean home range radii (as estimated by minimum convex polygons) is ∼ 8.5 km (Kolowski et al. 2010). We ran two models, a null model as well as a model that examined sex differences in detection. Forest elephant sex ratios are typically female biased (Fishlock and Lee 2013, Turkalo et al. 2013); thus, we modeled for this potential influencing factor in density estimation. While most samples yielded consistent genotypes at the three sex-determining loci, we classified the sex for any samples that did not consistently amplify as unknown. We compared these models using Akaike’s Information Criteria corrected for small sample size (AICc) and determined the best fit model using ΔAICc < 2 (Burnham and Anderson 2002).

### Sociality Analysis

We used fresh dung samples collected in close proximity to assess patterns of sociality. We excluded any samples that appeared to be of a different age based on appearance (moisture, degradation, etc.) than the surrounding samples (N = 4). Previous forest elephant studies have used varying distances to define association. Munshi-South (2011) used a distance of 100 m whereas Schuttler et al. (2014a,b) used 250 m. The latter was described by the authors to be a compromise between 50 m used in Morgan and Lee (2007) for forest elephants and 500 m used for savanna elephants in Wittemyer et al. (2005). As there is no consensus on the appropriate distance to define association in elephants and given that the distance used to define association will determine the subsequent social network, we constructed social networks using both 100 and 250 m as defining association. We also constructed a social network using 75 m to assess how different this social network would be compared to the 100 m and 250 m networks. Distances between samples were calculated in ArcMap, version 10.4 (ESRI 2015). Samples in association that were collected on the same day and estimated to have been deposited at the same time (based on the appearance of the sample) were considered to be individuals from the same social group (e.g., Brand et al. 2016).

In order to visualize the social organization of the individuals represented in our sample we used the package igraph (Csardi and Nepusz 2006) in R (R Core Team 2016). We coded individual vertices for sex using shape and for mitochondrial haplotype using color. Additionally, we weighted the edges of our social networks using estimated genetic relatedness calculated using ML-Relate (Kalinowski et al. 2006). We excluded individuals that were found in isolation from the network. We visualized networks both with males and without males and report the number of vertices (individuals), edges (associations between individuals), and components (groups of vertices) for each network.

In order to examine whether or not relatedness was correlated with association, we used our association data and Queller and Goodnight (1989) relatedness values calculated in GENALEX and ran Mantel tests in R using the package ‘vegan’ (Oksanen et al. 2019).

## Results

### Sampling, Individual Identification, and Sex

We visited 61 and 82 points in the dry and wet season, respectively, covering 84 of the 100 total points. We collected 300 total samples from 31 points. We calculated P(ID)_sib_ to be < 0.001 at 7 microsatellite loci. Given the high degree of sociality among forest elephants and need for more loci to distinguish among related individuals/genotypes, we excluded samples that could only be genotyped at fewer than 7 loci (N = 12). Our analysis yielded 288 samples after exclusion, constituting 190 unique individuals. Some elephants were recaptured at the same location and time (N = 53) so we excluded these ‘false recaptures’ for our abundance and density estimates but retained them for our sociality analyses. Recapture rates ranged from one to four (Table 2) and all recaptures matched the capture in sex and haplotype when either could be determined.

**Table 1.** Genetic diversity measures per locus. Na, number of alleles; Ne, number of effective alleles; Ho, Observed heterozygosity; He, expected heterozygosity; UHe, unbiased expected heterozygosity; F_IS_, Inbreeding coefficient; HWE, p-values for Hardy-Weinberg equilibrium test.

| Marker | Na | Ne | Ho | He | UHe | Fis | HWE |
| --- | --- | --- | --- | --- | --- | --- | --- |
| FH126 | 16 | 8.703 | 0.815 | 0.885 | 0.887 | 0.079 | NS |
| FH48R | 13 | 5.175 | 0.754 | 0.807 | 0.809 | 0.065 | < 0.05 |
| FH60R | 12 | 7.071 | 0.858 | 0.859 | 0.861 | 0.001 | NS |
| FH67 | 6 | 4.191 | 0.774 | 0.761 | 0.763 | -0.016 | NS |
| FH94R | 10 | 2.942 | 0.746 | 0.660 | 0.662 | -0.130 | NS |
| LA4 | 12 | 7.037 | 0.695 | 0.858 | 0.860 | 0.189 | < 0.001 |
| LA5 | 8 | 4.388 | 0.684 | 0.772 | 0.774 | 0.114 | < 0.01 |
| LA6R | 10 | 3.891 | 0.784 | 0.743 | 0.745 | -0.055 | NS |
| Mean | 10.875 | 5.425 | 0.764 | 0.793 | 0.795 | 0.031 |  |
| Std. Dev. | 3.09 | 1.97 | 0.06 | 0.07 | 0.07 | 0.10 |  |

**Table 2.** Capture rate of individual elephants.

| Capture(s) (N) | Individuals (N) | Females/Males/Unknowns (N) |
| --- | --- | --- |
| 1 | 157 | 84/65/8 |
| 2 | 25 | 12/10/3 |
| 3 | 5 | 2/1/2 |
| 4 | 2 | 0/2/0 |
| 5 | 1 | 1/0/0 |

### Genetic Diversity, Hardy-Weinberg Equilibrium, Linkage Disequilibrium, and Null Alleles

Genetic diversity measures from identified individuals are presented in Table 1, and Supplement 1 contains allelic dropout estimates and false allele rates. Our results are consistent with previous studies using these markers in forest elephants inhabiting the Gamba Complex Industrial Corridor, with the presence of loci out of HWE, loci in LD, and null alleles explained by population structure and non-random mating (Eggert et al. 2014, Johnson et al. 2019). Exclusion of the markers that may have contained null alleles did not change our results, including population size estimates, so we included them in our downstream analyses.

### Estimation of Abundance and Density

Confidence intervals for estimated population abundance varied across three Capwire models (Table 3) from 432 - 683 (ECM) to a high of 754 - 1502 (TIRMpart) (Table 3). These models were significantly different in how well each was supported: the TIRM was better supported than the ECM (likelihood ratio = 43.52, P < 0.05) and TIRMpart was better supported than the TIRM (P < 0.0001).

**Table 3.** Forest elephant abundance and density estimates per km^2^ in our study area

| **Abundance**<br>**Model (Capwire)** | **Model**<br>**Likelihood** | **Abundance**<br>**95% CI (Point Estimate)** | **Density**<br>**95% CI (Point Estimate)** |
| --- | --- | --- | --- |
| ECM | -1131.598 | 432 - 683 (530) |  |
| TIRM | -1088.083 | 604 - 1038 (690) |  |
| TIRMpart | -935.884 | 754 - 1502 (867) |  |
| **Density**<br>**Model (SECR)** |  |  |  |
| Null | -484.761 |  | 0.47 - 0.80 (0.62) |
| Sex Differences<br>in Detection<br>Probability | -596.839 |  | 0.49 - 0.84 (0.64) |

The null SECR model resulted in a population density of 0.62 elephants per km^2^ (95% CI: 0.47-0.80) whereas our SECR model examining sex differences in detection probability indicated 0.64 elephants per km^2^ (95% CI: 0.49 - 0.84). Based on AICc values, the first model better fit the data (ΔAICc = 230.49).

### Sociality

Overall, we found similarity between our three sets of social networks of varying association radii (75, 100, and 250 m) in that a substantial proportion of networks in all analyses contained females with different mitochondrial haplotypes. Thus, we here present data on the 100 m social networks. See Supplement 2 for results from the other networks. When including both sexes, the 100 m social networks consisted of 35 components, 166 edges, and 139 vertices (Figure 2a). The components ranged in size from 2 to 21 individuals. Excluding individuals of unknown sex, 11 components were composed of all females, 3 components were composed of all males, and 20 components consisted of both females and males. One component consisted of two individuals of unknown sex. We detected six mitochondrial haplotypes, all of which have been identified in previous studies (Johnson et al. 2007, Munshi-South 2011). Twenty-two components had more than one female. Twelve of these components (54.5%) exhibited the same mitochondrial haplotype. When males and individuals of unknown sex were excluded from the analysis, the 100 m social network comprised 24 components, 58 edges, and 66 vertices that ranged in size from 2 to 5 individuals (Figure 2b). We could not confirm the presence of multiple haplotypes in 3 components due to individuals lacking haplotype data. Of the 21 components we were able to assess, 12 (57%) had the same mitochondrial haplotype. Our other social networks with different association radii (75 m and 250 m) also included components with females that had different mitochondrial haplotypes whether or not males were included (Supplement 2).

**Figure 2.**
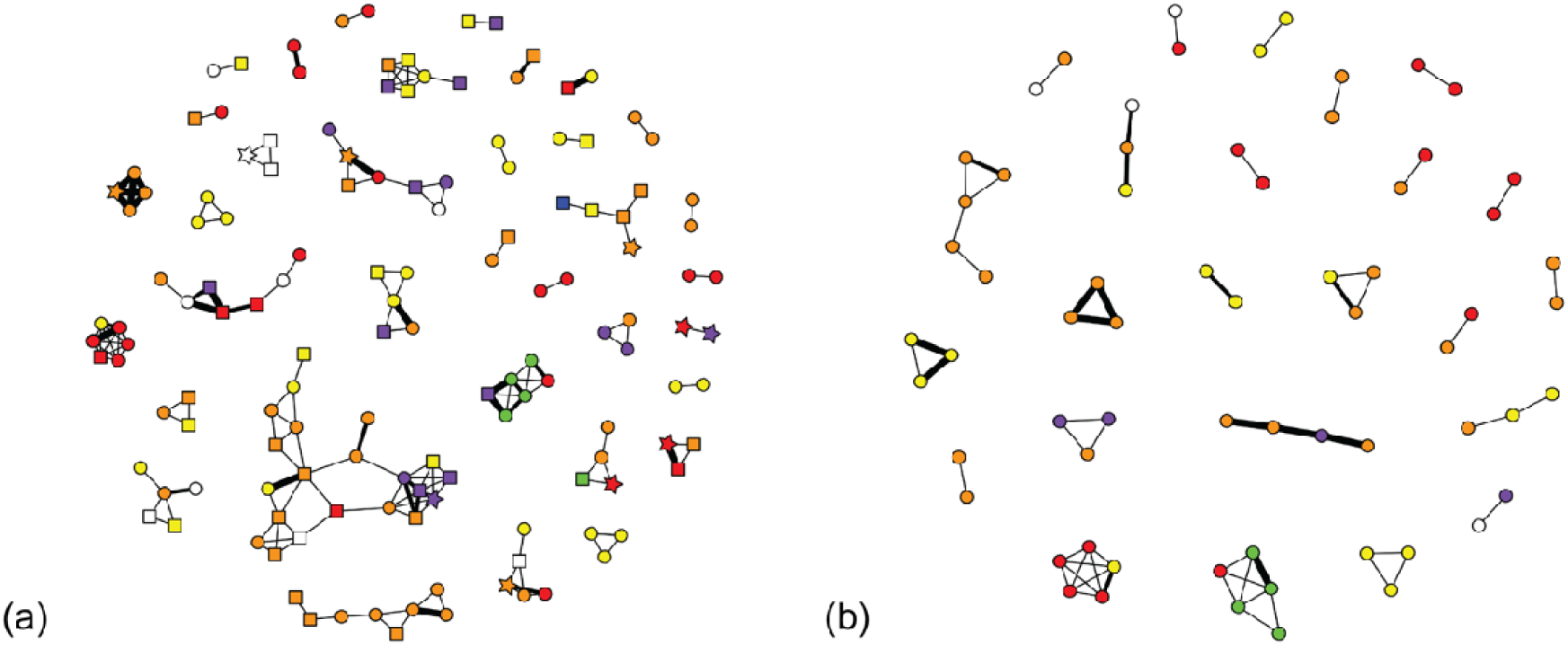
Social networks. The left network (2a) includes females (circles), males (squares), and individuals of unknown sex (stars) while the network on the right (2b) contains only females. Node color indicates mtDNA haplotype; white nodes are individuals whose mtDNA haplotype could not be determined. Edges are weighted by kinship estimated using ML-Relate with medium weighted lines representing second order relatives and the thickest lines representing first order relatives.

Mean relatedness across all sampled individuals was -0.006 ± 0.177. Among all dyads in all components in the 100 m social network, mean relatedness was 0.184 ± 0.021. When males and individuals of unknown sex were excluded, mean relatedness between all dyads in all components rose to 0.246 ± 0.033. Relatedness was significantly, positively correlated with being found in association within 100 m (r = 0.072, iterations = 999, p < 0.01).

## Discussion

Our analyses produced different yet overlapping population size estimates using the different Capwire models. Concordant with other genetic capture-recapture studies (e.g. Granjon et al. 2017), we found support for the TIRMpart model, which estimated that between 697 and 1,252 forest elephants inhabit our study area. This indicates that this population of forest elephants exhibits heterogeneity in capture probability. Differences in capture probability may reflect sex differences between males and females or may represent male movement in and out of the Industrial Corridor (Eggert et al. 2014). While our sampling period was relatively short (9 months) and similar to some sampling periods for other social mammals with slow life histories, such as great apes (e.g. McCarthy et al. 2015, Moore and Vigilant 2014), it is quite likely the population was not completely closed during this time period. In particular, individual elephant movement in and out of the study area may have reduced the likelihood of recapture, particularly because our sampling periods were in different seasons and at least some of the elephants in this population show seasonal movement (Eggert et al. 2014). Analyzing the two sampling sessions separately would also reduce likelihood of recapture, and recent studies have shown that longer sampling periods, despite increasing the likelihood of violating the closed population assumption, yield more precise and accurate population size estimates (Aranjelovic et al. 2010, Granjon et al. 2017). Regardless, violation of the closed population assumption and reduction in recapture probability likely artificially inflates abundance and density estimates, so that our results may actually be optimistic.

The best fit SECR model produced density estimates of 0.47 – 0.80 elephants per km^2^ that fall within the known range for forest elephants in Gabon: 0.18 elephants per km^2^ at Monts Birougou National Park to 1.06 elephants per km^2^ at Mwagné National Park (Turkalo et al. 2016). There is some evidence that elephants in the Gamba Complex are attracted to roads and other human infrastructure (H. Vanthomme, personal observation). If this is the case and given that sampling tended to be biased toward areas close to roads, these densities may represent the highest densities that can be found in the Gamba Complex.

If we extrapolate our results to the area of the entire Industrial Corridor (3,585 km^2^), we estimate that between 3,033 to 6,043 forest elephants inhabit this area based on abundance or between 1,684 to 2,832 elephants based on density. We note that extrapolation of these numbers assumes homogeneity in the factors that drive density across the landscape, which is unlikely. Since it is possible that human infrastructure may attract elephants, densities may be lower in other parts of the Industrial Corridor, which would make our extrapolated abundance estimates optimistic. Despite this variation, the extrapolated abundance estimates are still quite short of the estimate of 10,000 elephants in the Industrial Corridor (Eggert et al. 2014). This discrepancy may be explained by either differences in methodology and/or a recent decline in forest elephant population size in the Gamba Complex. Given the catastrophic population declines seen elsewhere in Gabon (Poulsen et al. 2017), this latter potential scenario is worrying. However, there is little evidence that elephant hunting and habitat loss in the Gamba Complex is at a rate that would cause such a rapid demographic decline. Still, new infrastructure as well as new mining and oil concessions pose a significant threat and underscore the need to preserve this area, especially given its importance as one of the last remaining forest elephant strongholds (Maisels et al. 2013).

We found that relatedness was consistent with our expectations from elephant social structure such that elephants found in association were more related than the mean relatedness across all individuals. Relatedness among associated females was also higher than the mean for all associated individuals. However, contrary to our prediction for forest elephant social structure based on previous studies and assumptions regarding the presence of strong female philopatry, we did not find a high proportion of social networks where females had the same mitochondrial haplotype. Approximately half of the networks exhibited females with only one mitochondrial haplotype, regardless of whether or not males and individuals of unknown sex were included in these networks (54.5% and 57%, respectively). Using a similar approach, a recent study examined social networks in forest elephants at Lopé (Schuttler et al. 2014a) and found that 79% of social network components that had multiple females shared the same mitochondrial haplotype, which is more concordant with the female philopatric social structure typically found in savanna elephants. This suggests that the forest elephants living in the southern region of the Industrial Corridor may exhibit some degree of female dispersal, which is different to what has been assumed and found for other forest elephant populations as well as most African savanna elephants. Johnson et al. (2019) found low but significant *F*_*ST*_ values in the mitochondrial DNA of these same animals. However, even modest levels of female dispersal could produce our observed pattern for social networks and relatedness. This departure from the expected pattern of female dispersal could be due to a number of reasons. It is possible that these social networks represent temporary fusion events. Alternatively, forest elephants may show considerable natural variation in social structure with conditional female dispersal as seen in other social mammals (e.g. Wikberg et al. 2012). In relation, local ecological conditions, such as the presence of plantations and/or high human population density, could be driving female dispersal and/or fusion events between different matrilines. As we excluded any samples that were differently aged, it is unlikely that we mistakenly assigned different matrilines feeding from the same resource at different times on the same day as being found in association. Further, multiple haplotypes appear to be more likely detected in closer proximity to human infrastructure, such as plantations. Lastly, it is also possible we are detecting anthropogenic effects such as agriculture or poaching on forest elephant behavior and social structure. For example, Johnson et al. (2019) recently reported on crop raiding in this forest elephant population, and poaching has been previously suggested to explain departures in expected social structure for savanna elephants in Mikumi National Park, Tanzania (Gobush et al. 2009), Queen Elizabeth National Park, Uganda (Nyakaana et al. 2001), and elsewhere (Archie and Chiyo 2012). As poaching disrupts social relationships between females, females left without a social group or smaller social groups may associate with other, unrelated females. Given the smaller size of forest elephant social units compared to savanna elephants (Munshi-South 2011, Schuttler et al. 2014), the death of even one female may result in an individual female without a social group. It’s possible that the human dominated landscape in Gamba may act as a refuge from poaching for elephants resulting in multiple sedentary matrilines. However, further research is required to fully understand the anthropogenic and ecological variables that drive variation in forest elephant social structure.

## Conclusions

This research provides novel baseline information on the forest elephants inhabiting the Gamba Complex. We believe the population size of forest elephants in the Gamba Complex Industrial Corridor, and thus the Gamba Complex of Protected Areas as a whole, to be much smaller (40 – 83% lower) than previously thought. Further, it is possible that our estimates of abundance and density are optimistic. This makes the future survival of forest elephants in general that much more precarious because the Gamba Complex is viewed as a forest elephant stronghold in Gabon, which in turn is seen as the stronghold for the global population of forest elephants. Further, we suggest that the female philopatric nature of forest elephants may be overstated in the current literature, and that forest elephants show a greater range of variation in social structure than previously thought. Lastly, given the difficulty in estimating forest elephant population sizes, we believe that this study provides promise for a standardized method for forest elephant censusing, and expanding and repeating this study is essential to monitor changes in forest elephant population size and sociality. It is clear that African forest elephants face significant threats from habitat loss, hunting and poaching, and other human related activities. Given the slow intrinsic growth rate of this particular taxon (Turkalo et al. 2016), continuous population decline will result in an increasingly difficult population recovery. It is imperative that known populations are monitored to provide accurate and precise data on the status of these populations and the global forest elephant population as a whole.

## Acknowledgments

We thank the Centre National de la Recherche Scientifique et Technologique for authorizing our study (permits #AR0017/13/MENESTFPRSCJS/CENAREST/CG/CST/CSAR). We also thank the Gabon-Oregon Center, the Smithsonian Conservation Biology Institute, and Shell Gabon for funding, and Rob Fleischer and Nancy McInerney for all their help and support for this project at the Center for Conservation Genomics at the National Zoological Park. Many thanks to our collaborators who contributed to field work: M. Onbenotori, G. R. Mihindou, A. Litona-Boubeya, and N. Simons. We also thank Murray Efford who graciously helped with the SECR analyses. This is contribution #192 of the Gabon Biodiversity Program.

## Supplements

1. Allelic dropout and false allele rate per locus.
2. Additional social networks.

