## Supplement 1 for "Abundance, density, and social structure of African forest elephants (*Loxodonta cyclotis*) in a human-modified landscape in southwestern Gabon"

Supplement 1. Allelic dropout and false allele rate per locus.

| Marker | Dropout | False Allele |
| --- | --- | --- |
| FH126 | 0.032 | 0.00 |
| FH48R | 0.043 | 0.00 |
| FH60R | 0.079 | 0.00 |
| FH67 | 0.011 | 0.00 |
| FH94R | 0.000 | 0.34 |
| LA4 | 0.087 | 0.00 |
| LA5 | 0.169 | 0.00 |
| LA6R | 0.180 | 0.00 |
