## Supplement 2 for "Abundance, density, and social structure of African forest elephants (*Loxodonta cyclotis*) in a human-modified landscape in southwestern Gabon"

Supplement 2. Additional social networks.

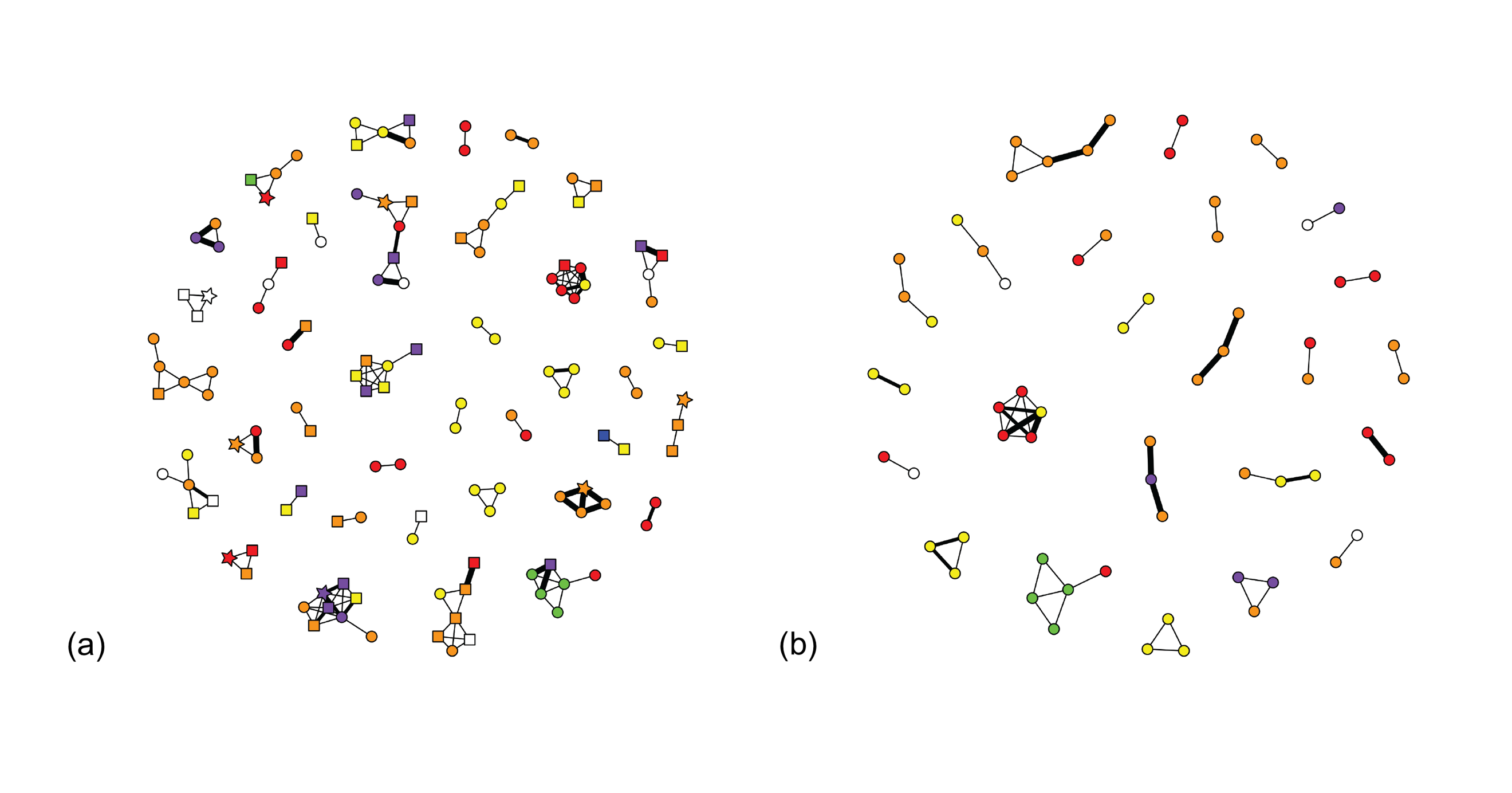

Figure 1. 75 m radius social network. The left network (1a) includes females (circles), males (squares), and individuals of unknown sex (stars) while the network on the right (1b) contains only females. Node color indicates mtDNA haplotype; white nodes are individuals whose mtDNA haplotype could not be determined. Edges are weighted by kinship estimated using ML-Relate with medium weighted lines representing second order relatives and the thickest lines representing first order relatives. 1a has 132 vertices, 148 edges, and 38 components whereas 1b has 65 vertices, 53 edges, and 23 components

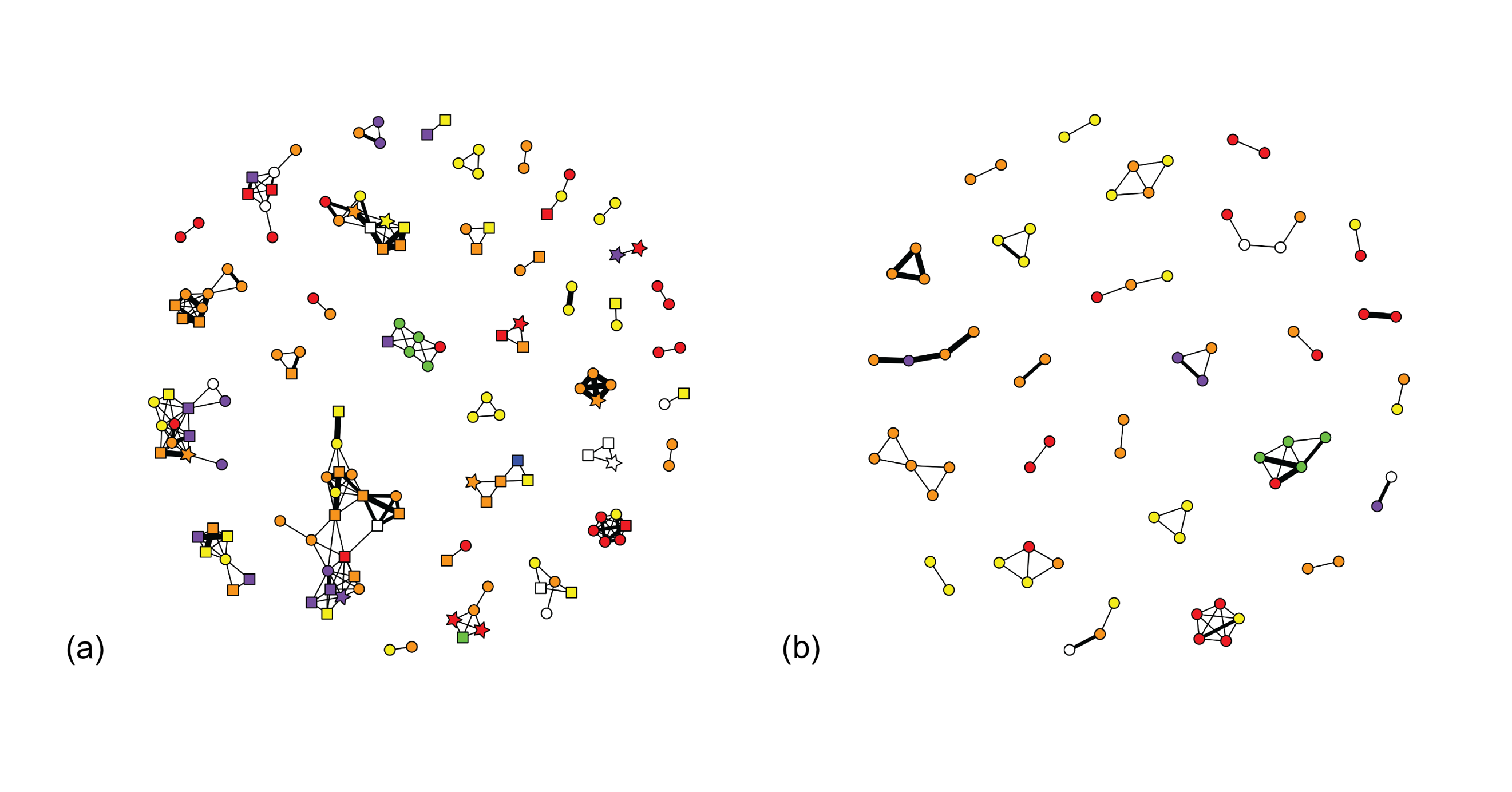

Figure 2. 250 m radius social network. The left network (2a) includes females (circles), males (squares), and individuals of unknown sex (stars) while the network on the right (2b) contains only females. Node color indicates mtDNA haplotype; white nodes are individuals whose mtDNA haplotype could not be determined. Edges are weighted by kinship estimated using ML-Relate with medium weighted lines representing second order relatives and the thickest lines representing first order relatives. relatives. 2a has 149 vertices, 235 edges, and 35 components whereas 2b has 75 vertices, 69 edges, and 26 components.
